# Integrating somatosensory information over time

**DOI:** 10.1101/817262

**Authors:** Raúl Hernández-Pérez, Eduardo Rojas-Hortelano, Victor de Lafuente

**Affiliations:** Institute of Neurobiology, National Autonomous University of Mexico. Boulevard Juriquilla 3001, Querétaro, QRO., México, 76230

**Keywords:** touch, somatosensory, accumulation-to-bound, independent sampling, decision-making.

## Abstract

Our choices are often informed by temporally integrating streams of sensory information. This has been well demonstrated in the visual and auditory domains, but the integration of tactile information over time has been less studied. We designed an active touch task in which subjects explored a spheroid-shaped object to determine its inclination with respect to the horizontal plane (inclined to the left or to the right). In agreement with previous findings, our results show that more errors, and longer decision times, accompany difficult decisions (small inclination angles). To gain insight into the decision-making process, we used a task in which the time available for tactile exploration was varied by the experimenter, in a trial-by-trial basis. The behavioral results were fit with a model of bounded accumulation, and also with an *independent-sampling* model which assumes no sensory accumulation. The results of model fits favor an *accumulation-to-bound* mechanism, and suggest that participants integrate the first 600 ms of 1800 ms-long stimuli. This means that the somatosensory system benefits from longer streams of information although it does not make use of all available evidence.

**Highlights:**

- The somatosensory system integrates information streams through time.
- Somatosensory discrimination thresholds decrease with longer stimuli.
- A bounded accumulation model is favored over independent sampling.
- Humans accumulate up to 600 ms, out of 1800 ms-long stimuli.

## Introduction

A fundamental aim in systems neuroscience is elucidating the mechanisms underlying behavioral decisions, which are often based on noisy or ambiguous sensory information (Andersen & Cui, 2009; Romo & de Lafuente, 2013; Shadlen & Kiani, 2013; Hanks & Summerfield, 2017). A core behavioral observation in decision-making experiments is that difficult decisions lead to more errors and, importantly, take more time to complete. This behavioral pattern has been fruitfully studied under the quantitative framework of an *accumulation-to-bound* decision model (Bogacz, Brown, Moehlis, Holmes, & Cohen, 2006; Kiani, Hanks, & Shadlen, 2008; Ratcliff, Smith, Brown, & McKoon, 2016; Tsetsos, Gao, McClelland, & Usher, 2012). The temporal integration of sensory streams up to a decision bound that favors a particular choice, simultaneously explains why more errors are made when information is noisy, and why these decisions take more time to complete. Examples of the success of the *accumulation-to-bound* model to explain behavioral performance and neuronal activity constitute a rich literature (Huk & Shadlen, 2005; Donner, Siegel, Fries, & Engel, 2009; de Lafuente, Jazayeri, & Shadlen, 2015; Brunton, Botvinick, & Brody, 2013; Cisek & Kalaska, 2010; Ding & Gold, 2013; Bollimunta, Totten, & Ditterich, 2012; Heekeren, Marrett, & Ungerleider, 2008). However, in the majority of those experiments, subjects were trained to use visual or auditory information to inform their decisions.

Two recent studies have addressed accumulation of somatosensory information. Delis et. al. (Delis, Dmochowski, Sajda, & Wang, 2018) found that the behavior of human subjects in an active surface texture discrimination, and the underlying EEG signals, were compatible with a process of accumulation of somatosensory information occurring in the middle frontal gyrus (see also Pleger et al., 2006)). Nest et. al. (2017) demonstrated that the discrimination threshold for the amplitude of body motion (self-motion) decreased with increasing stimulus duration, and this threshold decrease was compatible with a process of leaky integration of vestibular sensory information. These two important studies suggest that *accumulation-to-bound* of somatosensory information is a plausible decision mechanism that is compatible with behavioral data and somatosensory neuronal activity (see also (Drugowitsch, DeAngelis, Klier, Angelaki, & Pouget, 2014; Mulder & van Maanen, 2013)). However, to our knowledge ours is the first study to test whether a mechanism of *accumulation-to-bound*, or instead, a simpler *independent sampling* model better explains the decrease in threshold observed for longer sensory stimuli.

Our experimental approach allowed to test whether an alternative model that assumes no temporal integration, could explain the core observation that ambiguous information lead to more errors and longer decision times. This model, often termed *probability summation* or *independent sampling* (Palmer, Huk, & Shadlen, 2005; Wandell, Ahumada, & Welsh, 1984; Watson, 1979) proposes that streams of sensory information are not accumulated, but sampled independently to determine if the most recent piece of evidence is large enough to reach a decision bound. Testing alternative models of decision-making in the somatosensory system will allow establishing which information-processing mechanisms might be shared among sensory systems (Herding, Ludwig, von Lautz, Spitzer, & Blankenburg, 2019; O’Connell, Dockree, & Kelly, 2012; Pack & Bensmaia, 2015; Yau, Pasupathy, Fitzgerald, Hsiao, & Connor, 2009).

We performed active tactile decision-making experiments in which subjects had to touch an object to determine its inclination. The spheroid-shaped object that participants explored could be inclined clockwise or counter-clockwise with respect the horizontal plane. Our results show that the somatosensory system is indeed capable of temporally integrating streams of sensory evidence. However, perfect accumulation of information was achieved for only a third of the total duration of the stimulus.

## Experimental procedures

### Reaction-time orientation discrimination task

Subjects performed an active tactile task in which they had to touch an elliptical object to determine whether its inclination was clockwise (*rightward*) or counterclockwise (*leftward*) with respect to the horizontal plane (Figure 1; n=8 participants, 4 female, mean age 25, range 18-31). Each trial began with an auditory cue that instructed the subject to touch the object (with their right hand) to determine its inclination. They wore a bracelet that closed an electrical circuit when the hand made contact with the object (voltage sampled at 1 KHz, Measurement Computing USB-1208 FS, Middleboro, MA, USA). After exploring the object and committing to a choice, participants released the object and communicated their decision with the left hand by means of a computer mouse (right click for *rightward* rotations, left click for *leftward* rotations). They were given no specific instructions as to how much time they should take to make their decisions, thus they were free to take as much time as needed. This task design defines what is called a *reaction time task*. Correct and incorrect trials were signaled to the subjects by means of auditory cues.

**Figure 1.**
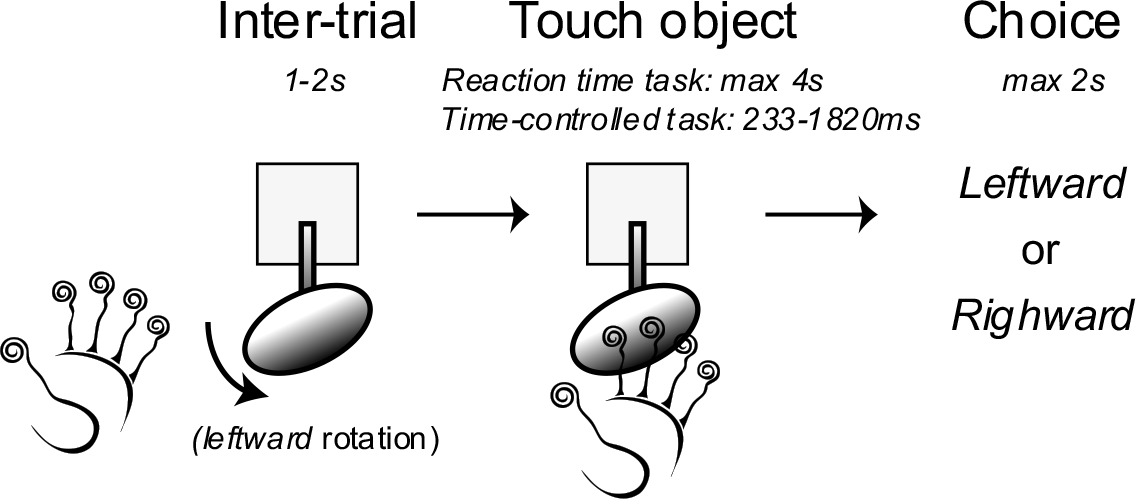
Orientation-discrimination task. After an inter-trial interval (1-2s) participants received an auditory cue that instructed them to touch the object to determine whether its inclination was *leftward* of *rightward* with respect to the horizontal plane. In the *reaction-time* task subjects had up to 4s to explore the object and communicate their choice. In the *time-controlled* task, exploration time was sampled from an exponential distribution (233-1820 ms). Once the subjects released the object, they had up to 2s to communicate their choice. Feedback for correct and incorrect trials was provided by auditory cues immediately after the mouse click that indicated their choice (using their left hand; eyes were covered, see Methods).

Participants seated comfortably in a quiet room with their eyes covered and wore headphones with white noise that also presented the auditory instructions for the task. They completed the task in four 15-min blocks (with rest periods interleaved), performing 144 trials per stimulus orientation (10 orientations) for a total of 1440 trials per subject. Written consent was obtained prior to the study, and subjects were free to abandon the study at any time. They were compensated monetarily for their participation. Procedures were in accordance with the Declaration of Helsinki and were approved by the Institute for Neurobiology’s Ethics Committee. The object was made from machined aluminum and had the shape of a spheroid (similar to an oval-shaped football). It measured 13 cm in its long axis and 2.5 cm in the two orthogonal axes. The direction and the angle of inclination were pseudo-randomly selected and were controlled by a servo motor under Matlab control (Hitec, HS-815BB Mega Sail Servo, Poway, CA, USA). It is important to note that rotations of the object occurred during the inter-trial interval, not while subjects were touching the object.

### Time-controlled orientation discrimination task

The second task, that we termed the *time-controlled*, progressed just as described, but instead of subjects moving their hand to touch the object, the object moved towards and away from the subjects’ hand (13 participants, 7 female, 18-32 age range, 25.2 mean age). With the elbow resting and comfortably fixed to the table with Velcro straps, subjects opened their hands as if to catch a ball to wait for the stimulus. The object traveled forward 15 cm (on wheels over a rail) to make contact with the subjects’ hand. It traveled backwards the same 15 cm to remove the object from the hand. The duration of contact between the hand and the object (stimulus duration) was randomly selected from a truncated exponential distribution, ranging from 233 to 1820 ms. This exponential distribution of stimulus durations was divided into 10 bins of increasing widths so that each bin had the same number of trials. For each duration bin, we used a staircase procedure to calculate the orientation discrimination threshold. In the 2-up 1-down staircase procedure we used, the stimulus amplitude (inclination angle) increased after an error and decreased after two correct responses. A change in direction, from a stimulus increase to decrease, or from decrease to increase, was labeled as a reversal. The last 13 of 15 reversals were used to estimate a discrimination threshold (angle) corresponding to 80.35% correct responses (Kingdom & Prins, 2010).

### Independent sampling and Accumulation-to-bound models

To gain insight into the mechanism by which a stream of sensory evidence is transformed into a decision we tested two models (Figure 2). The models assume there is a *decision variable* that represents sensory evidence, moment-by-moment in *independent sampling*, and summed through time in *accumulation-to-bound*. The probability distributions of the *decision variable* for the two models are illustrated in Figure 2 (three time steps are shown). The mean of the distributions is determined by the stimulus angle, and is defined as the product *k·stimulusAngle*, where *k* is a sensitivity parameter. The standard deviation (*std*) of the distribution at time zero is one, so *A* and *k* can be thought of as having *std* units. Note how the probability distribution in the *independent sampling* model is constant through time (Figure 2B), and note how in the *accumulation-to-bound* model the mean and *std* increase as a function of time due to the accumulation of sensory evidence (Figure 2A). To simulate the evolution of the *decision variable* in the *accumulation-to-bound* model, on each time step the distribution is convolved with the initial gaussian distribution.

**Figure 2.**
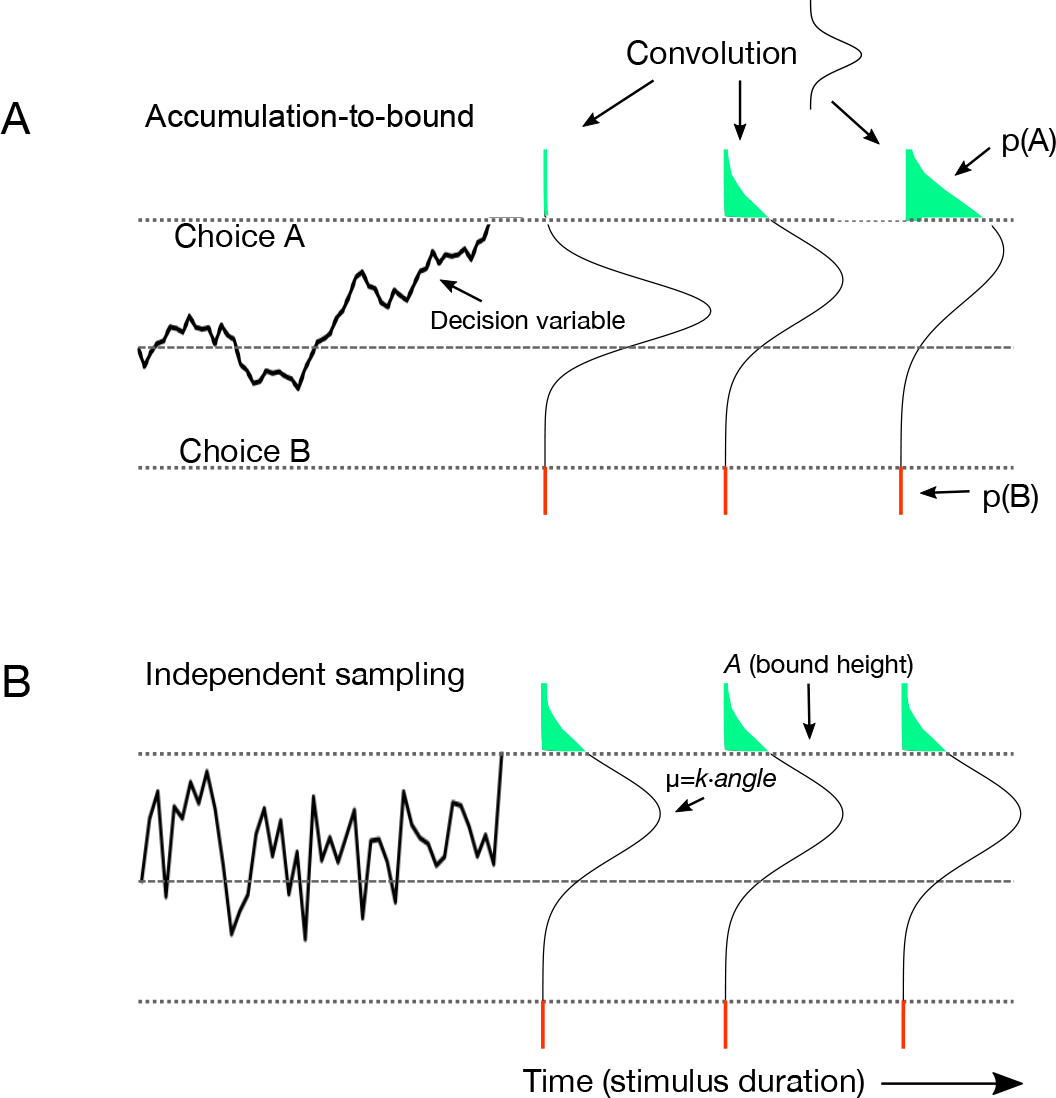
Accumulation-to-bound and Independent sampling models. (**A**) In *accumulation-to-bound*, the probability distribution of the decision variable representing the accumulated sensory evidence was modeled by a gaussian distribution. The mean of this distribution was determined by the product of the stimulus angle and the sensitivity parameter *k*. The sign of the angle thus determines whether the mean of the decision variable drifts towards Choice A or B. To simulate this drift, on each time step, the distribution is recursively convolved with the distribution at time zero. Then, the probabilities p(A) and p(B) are used to calculate the probability of selecting Choice A (see Methods). (**B**) The only difference in the *independent sampling* model is that sensory evidence does not accumulate. In the simulation, the distribution of the decision variable remains constant through time (no convolution operation).

To calculate the probability of a correct response for the *independent sampling* model we used the formula:

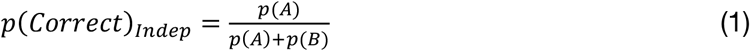

where *p(A)* and *p(B)* are the probabilities of the decision variable reaching the upper and lower bounds, respectively (Figure 2). We designated the upper bound to be the correct bound. To fit reaction time data, we used:

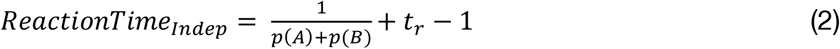

where *t*_*r*_ is the residual time attributable to non-decision processes such as motor movement initiation.

For the accumulation-to-bound we used the functions (Palmer et al., 2005):

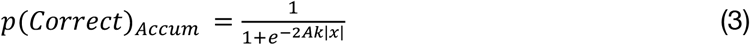

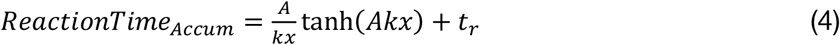

For each model, we simultaneously fit the *p(Correct)* and *reaction time* data using the non-linear least squares method of Matlab’s *Fit* function. The fitted parameters were: bound height (*A*), sensitivity (*k*), and residual time (*t*_*r*_).

Regarding the results of the second experiment (*time-controlled* task), to calculate *p(Correct)* as a function of stimulus duration for the *independent sampling* model we used the equation:

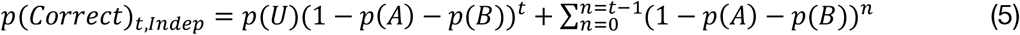

In *accumulation-to-bound p(Correct)*_*t*_ is also calculated with this formula, but on each time step the probability distribution is convolved with a gaussian kernel to simulate the evolving dynamics of the accumulated *decision variable* (Figure 2). For both models, we used Matlab’s *fmincon* function to fit the parameters: bound height (*A*), sensitivity (*k*), and a location parameter for the *y-axis* that sums to the threshold.

Figure 3 illustrates how the models predict the decrease in discrimination threshold as a function of stimulus duration. In *accumulation-to-bound*, the threshold first decreases with a −½ slope, indicating that there is perfect accumulation. As stimulus duration increases, the threshold eventually flattens and there are no further gains from longer stimulus durations. The decrease in threshold flattens because, once the decision variable reaches a bound, it can no longer integrate further evidence. The decrease in threshold for *independent sampling* does not show a −½ slope, but instead predicts a sigmoidal decrease of the threshold that eventually flattens at the same level as *accumulation-to-bound*, although for much longer stimulus durations. These two models were fit to the *time-controlled* tactile task data described earlier.

**Figure 3.**
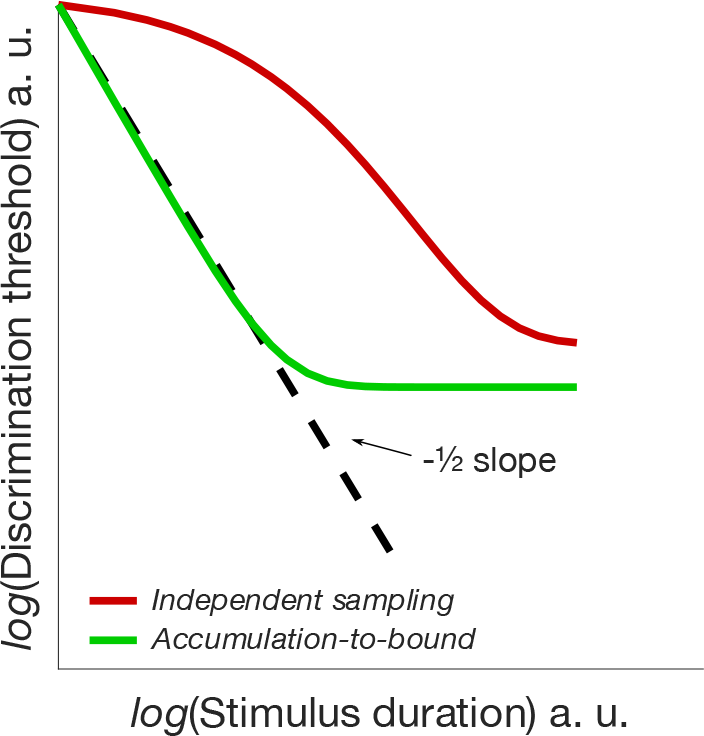
Model predictions. The *accumulation-to-bound* model predicts that the discrimination threshold should decrease with a −½ slope for short duration stimuli. The discrimination threshold eventually flattens due to the bound stopping the accumulation of sensory evidence, preventing further gains in sensibility. In the *independent sampling* model, the decrease in threshold never reaches a −½ slope and instead has a sigmoidal shape that eventually reaches the bounded accumulation threshold, although for much longer stimulus durations. The parameters of these two models were fit to the behavioral data obtained in the *time-controlled* experiment (see Methods and Results).

## Results

### Reaction-time task

In our first experiment, subjects touched an object (eyes covered) with their right hand to determine if its inclination was *rightward* (clockwise) or *leftward* (counterclockwise) with respect of the horizontal plane (Figure 1). On each inter-trial interval, the spheroid-shaped object was rotated to one of ±5 possible angles ranging from 2 to 32 degrees. After an auditory cue, subjects explored the object and took as much time as needed (up to 4s) to communicate their choice by clicking the right or left button of a computer mouse.

Figure 4A depicts how the probability of a *rightward* choice changes as a function of object inclination. The *psychometric* curve demonstrates that participants (n=8, see Methods) correctly performed the orientation discrimination task, rarely making mistakes at large angles, and displaying mixed responses at intermediate and small angles.

**Figure 4.**
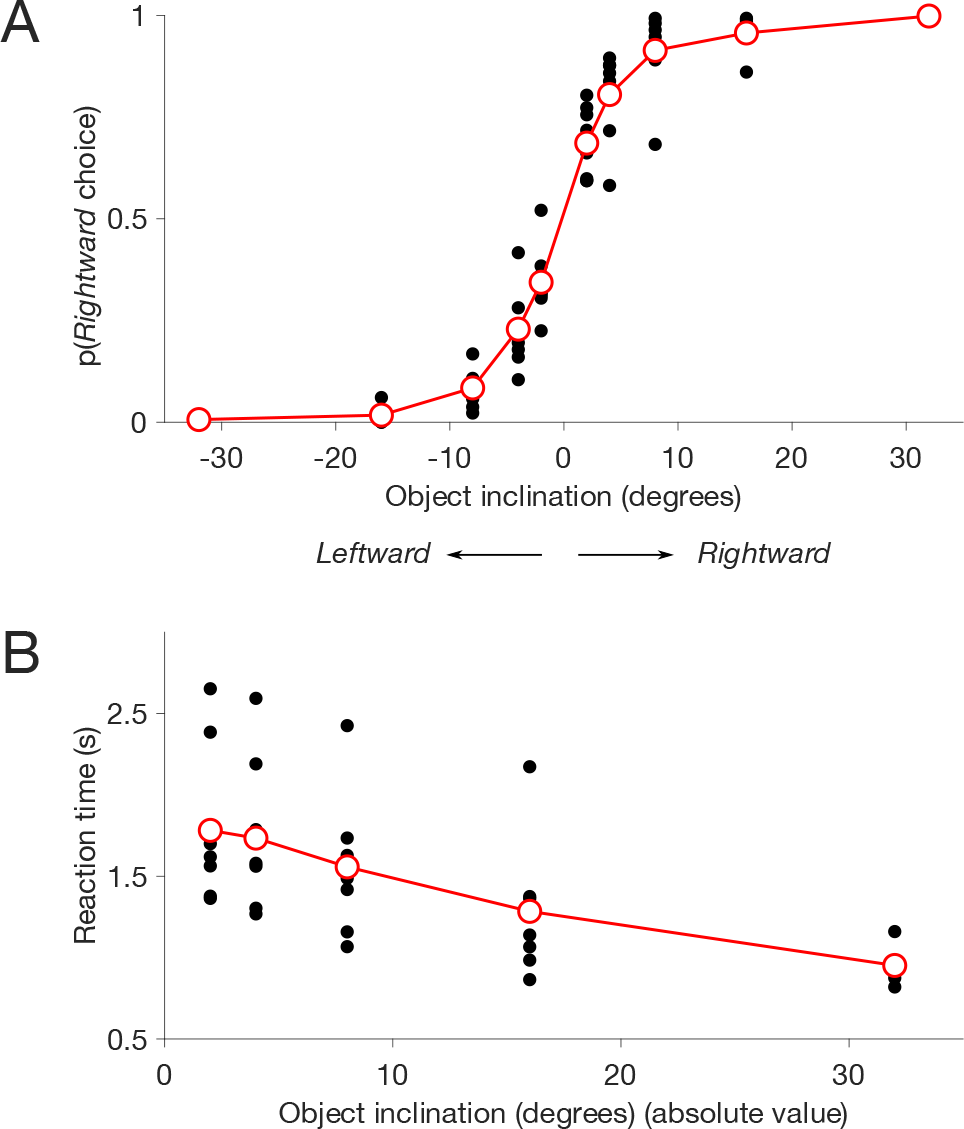
Psychometric and chronometric curves. (**A**) The probability of a *rightward* choice is plotted as a function of object inclination (degrees). A positive angle indicates clockwise inclination (*rightward*). Dots depict mean choice of 144 trials for each object angle, for each participant (n=8 subjects, 10 angles). Note the sigmoidal transition from perfect *leftward* to perfect *rightward* choices. Red line and markers indicate mean across participants. (**B**) Reaction time as a function of object inclination. Dots indicate the mean reaction time for each subject (n=8) and each inclination angle (absolute value of inclination angle, 244 trials per dot). The red line and markers indicate the mean across participants. Note how reaction times decrease with larger inclination angles.

The time subjects took to complete their decisions, as a function of object inclination, constitutes the *chronometric* curve, which can be seen in Figure 4B. It shows that small inclination angles result in longer exploration time by the subjects. This crucial characteristic of the *chronometric* curve is consistent with a model in which a *decision variable* reaches a bound to trigger a choice: large inclination angles quickly take the *decision variable* towards the decision bound, rapidly determining the behavioral choice. However, as we will demonstrate next, it does not necessarily validate a model of *bounded accumulation* of sensory evidence.

### Accumulation and Independent sampling account for reaction time and accuracy

An alternative to the *accumulation-to-bound* model is one that has been called *probability summation*, or *independent sampling*, and proposes that no accumulation takes place. Instead, decisions are reached when a spike (salient peak of sensory information), is large enough to reach one of the decision bounds. This is what is called *independent sampling* of the stream of sensory information. We wanted to explore if this alternative model is consistent with the *psychometric* and *chronometric* curves obtained in the *reaction-time* task.

Figure 5 shows the fits of the *independent sampling* and the *accumulation-to-bound* models to the behavioral data. The results indicate that both models provide satisfactory fits to the data. Both models are consistent with the *psychometric* and *chronometric* curves obtained from the *reaction-time* tactile discrimination task. These results indicate that the observation of increasing decision times with increasingly ambiguous sensory evidence, does not necessarily favor an *accumulation-to-bound* model of decision-making.

**Figure 5.**
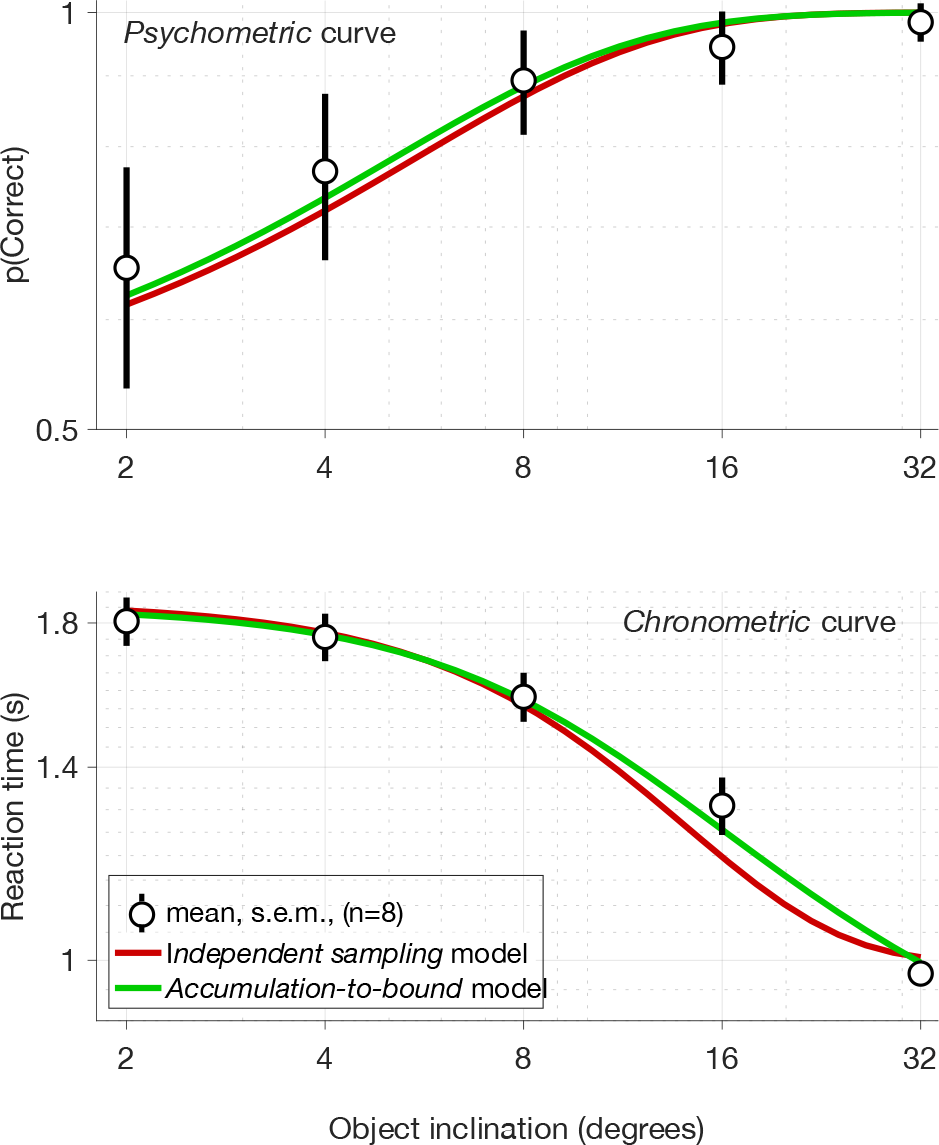
Model fits to accuracy and reaction time. The probability of a correct response is plotted as a function of object angle in the upper panel. Black circles and lines denote mean and s.e.m of *p(Correct)* across subjects (n=8). The logarithm of the absolute value of stimulus angle is used for the x-axis (same data as that in Figure 5A). The black circles and lines in the lower panel depict mean and s.e.m of the across-subject reaction times. Red and green lines are simultaneous fits to accuracy and reaction time of the *independent sampling* and *accumulation-to-bound* models, respectively. Note how both models provide satisfactory fits to the data.

Thus, we conclude that a *reaction time* experiment, in which subjects control the duration of the tactile exploration, did not allow establishing whether subjects accumulate information, or instead, they take independent samples of the sensory stream until a salient peak of information takes the decision variable to the decision bound. For this reason, we designed a new behavioral task in which the time available for exploring the object was controlled by the experimenter.

### Time-controlled task

The *accumulation-to-bound* model predicts that the discrimination threshold decreases as function of stimulus duration (Kiani et al., 2008). Moreover, perfect accumulation predicts that the decrease in threshold, when plotted in a log-log graph of threshold *vs* stimulus duration, has a −½ slope (Figure 3). It also predicts that the slope eventually flattens due the bound effectively stopping evidence accumulation, precluding further gains in sensitivity.

We wanted to test if the −½ slope and the flatting of the discrimination threshold, as precited by the *accumulation-to-bound* model, was present in the behavior of the subjects performing a tactile discrimination task. For this, we developed a version of the tactile task in which the duration of the contact of the hand with the object was varied by the experimenter (see Methods). The tactile object was mounted in a movable platform that allowed control of the touch duration, ranging from a few hundred milliseconds, up to several seconds. This new version of the task allowed us to estimate the increase in sensibility (i.e. decrease in discrimination threshold) resulting from longer stimulus durations. If subjects have the ability to perfectly accumulate tactile information, as has been demonstrated in other sensory systems, we would have obtained a line with a −½ slope relating discrimination threshold to stimulus duration.

The across-participants mean discrimination thresholds, plotted as a function of stimulus duration, is shown in Figure 6. Inspection of the data (black circles) suggest that subjects do not display perfect integration for the total duration of the stimulus. This can be seen as the departure of the data from the broken black line illustrating a −½ slope. This departure occurs at a stimulus duration of ~600 ms.

**Figure 6.**
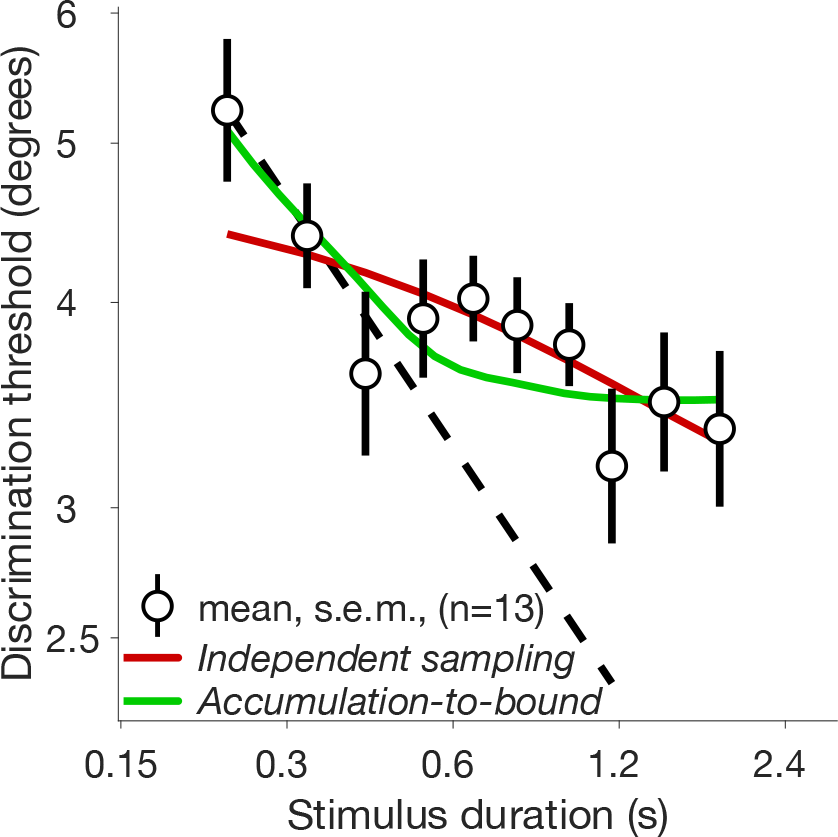
Decrease in threshold obtained from longer stimulus durations. Mean discrimination threshold (n=13 participants) is plotted as a function of stimulus duration. Black markers and vertical lines denote mean and s.e.m., respectively. The dashed line has a −½ slope, for reference. The accumulation-to-bound and independent sampling model fits to the data are plotted in green and red, respectively. Goodness of fit statistics indicate that the accumulation-to-bound model provides a better description of the behavioral data.

To quantify this observation we first carried, for each participant, a linear regression of the discrimination threshold as a function of stimulus duration (individual data and regressions not shown). Across subjects, the mean slope of improvement was −0.15 (±0.05 s.e.m), a much shallower slope than the −½ slope predicted by perfect accumulation.

Having discarded perfect accumulation for the total duration of the stimulus, we next aimed to specifically test which of the two models (*accumulation-to-bound* or *independent sampling*) better explained the decrease in threshold obtained from longer stimulus durations. First, we performed fits of the two models to the data of each participant. The results demonstrated that both models provided satisfactory fits to individual data. This was quantified by calculating the correlation coefficient between the discrimination thresholds predicted by the model, and that actually measured from the subjects (one correlation coefficient for each model and for each subject). A fisher transformed t-test on the fisher-corrected correlation coefficients confirmed that the difference between the model fits was not significant. That is, both models provided equally satisfactory descriptions of individual data (p=0.76; fits to individual data not shown).

Finally, we performed fits of the two models to the across-subjects pooled discrimination thresholds (Figure 6). The results show that both models provide satisfactory fits to the data in the sense that model predictions and measured thresholds are significant correlated (pearson correlation coefficient, p<0.01); and also predictions and data could not be distinguished by a two-sample Kolmogorov-Smirnov test (p>0.05 for both models). However, goodness of fit statistics indicate that the *accumulation-to-bound* model provides a better description of the data. The sum of squares due to error (*SSE*), and the root mean squared error (*RMSE*) both are lower for the *accumulation-to-bound* model (*SSE*_*accum*_=0.04, *SSE*_*indep*_=0.07; *RMSE*_*accum*_=0.09, *RSME*_*indep*_=0.11). Additionally, the coefficient of determination (*R^2^*) is larger for the *accumulation-to-bound* model (*R^2^*_*accum*_=0.76, *R^2^*_*indep*_=0.45). These goodness-of-fit statistics provide support to the hypothesis that there is bounded accumulation of somatosensory information, and that the bound stops accumulation process at approximately 600 ms after stimulus onset. Thus, our results show that only the first third of an 1800 ms-long stimulus is integrated to make a somatosensory decision.

## Discussion

The results of our first experiment, in which the participants determined exploration time (*reaction time* experiment), demonstrated that increasing reaction times as a function of decision difficulty do not necessarily imply that a*ccumulation-to-bound* should be favored over and *independent sampling* model. To disambiguate these models, we showed that it was necessary to perform a psychophysical task in which exploration time is controlled by the experimenter. In the second experiment, by systematically varying exploration time, we were able to estimate how the inclination discrimination threshold diminished as a function of stimulus duration. The results of this *time-controlled* tactile task provided support in favor of the idea that somatosensory information is accumulated for ~600 ms, from an 1800 ms sensory stream. Our results demonstrate that the *accumulation-to-bound* mechanism for transforming streams of sensory evidence into decisions is shared at least between the visual, auditory, and somatosensory systems.

Previous studies have suggested that by changing an integration time constant, a leaky integrator can be adapted to match the expected duration of stimuli (Ossmy et al., 2013; Waskom & Kiani, 2018). In monkeys, the decrease in the detection and discrimination thresholds for intra-cortical microstimulation stimuli have been shown to flatten at 200 and 300 ms, respectively (Kim et al., 2015). Thus, it is likely that the 600 ms integration period that we found can be modified by varying the mean expected duration of stimuli, so that shorted mean durations would yield shorter integration time windows. Our models did not include a leak parameter, and we only tested the extremes of leaky integration, i.e. no leak in perfect accumulation, and no accumulation in independent sampling. A leak parameter would certainly improve the fits to our behavioral data, although it has been shown that perfect bounded accumulation accounts for the trial-by-trial responses (Brunton et al., 2013) and also the decrease in threshold even for very long stimuli (>10 s; Waskom & Kiani, 2018).

We think it is important to note that our tasks were active exploration tasks which required subjects to actively explore the object instead of passively wait for the stimulus. This active exploration approach, engaging tactile and proprioceptive receptors as well as reaching and grasping motor plans (Alvarez, Zainos, & Romo, 2015; Callier, Suresh, & Bensmaia, 2019; Coallier & Kalaska, 2014; Delhaye, Xia, & Bensmaia, 2019; Kuang, Morel, & Gail, 2015; Michaels, Dann, Intveld, & Scherberger, 2018; Scherberger & Andersen, 2007), might have influenced our results in the sense that touching an object to determine its orientation might favor a shallow bound, precluding the perfect information accumulation that has been observed other contexts in which subjects passively wait for sensory information to arrive. The fact that subjects reached almost perfect levels of performance at high rotation angles (Figure 4A) indicates that subjects knew the task well enough. However, we think that longer training times in the *time-controlled* task time might help improve the ability to accumulate sensory information for longer time windows. We think that contact force, which we did not control, might be a particularly important variable to monitor in the future.

Our results showed that the *accumulation-to-bound* model was favored by the goodness-of-fit statistics of the model to the across-participants pooled data. However, when we fitted the model to each participant, we failed to confirm the hypothesis that *accumulation-to-bound* provided a better description of to the decrease in threshold for longer stimulus durations. This means that data for each participant was highly variable, and did not provided enough statistical power to disambiguate the models for individual results. In the future, this could be improved by increasing the number of training sessions, and the number of trials that each participant performs.

## Conclusion

Human subjects integrate somatosensory information streams (tactile and proprioceptive) arising from actively touching an object to determine its spatial orientation. The *accumulation-to-bound* model provides a good description of decision-making behavior in at least the visual, auditory, and somatosensory domains. The somatosensory system is able to benefit from longer stimulus streams but does not show perfect integration beyond ~600 ms of 1800 ms long stimuli. The decrease in discrimination threshold as a function of stimulus duration must be evaluated to disambiguate *accumulation-to-bound* from *independent sampling* models.

## Acknowledgements

We thank Edgar Bolaños for technical assistance. This research was supported by Dirección del Personal Académico de la Universidad Nacional Autónoma de México (PAPIIT IN207818) and CONACYT (VdL: 254313, 247200).

